# Insulotaxy: Navigating the Human Insula with a Novel Stereotactic Framework

**DOI:** 10.64898/2026.08.08.739317

**Authors:** Panagiotis Kerezoudis, Michael A Jensen, Bryan T. Klassen, Gregory A. Worrell, Nicholas M. Gregg, Nuri F. Ince, Jamie J. Van Gompel, Kai J. Miller

## Abstract

**Introduction:** The insula is an increasingly important target for functional neurosurgery given its involvement in a range of neurological and neuropsychiatric disorders, including epilepsy and chronic pain. As this practice evolves, optimal targeting will require standardized outcome measures that relate electrode or laser trajectory to postprocedural outcome. Traditional whole-brain registration approaches fail to capture the substantial person-to-person variability in insular gyral configuration, including the relative internal rotation of the insular gyri with respect to standard stereotactic space.

**Objective:** We propose and validate a stereotactic coordinate system based on local anatomical landmarks to facilitate surgical planning and standardized outcome assessment within the insular cortex.

**Methods:** Our approach transforms brain MRI first into standard AC-PC space, and then into an insular-specific space defined by five anatomical landmarks: four points along the central sulcus of the insula and one point at the middle cerebral artery (MCA) bifurcation (at the limen insulae). The system calculates two angles - θ (axial) and φ (sagittal) - between the AC-PC line and the insular axis, and the brain volume undergoes sequential rotation through these angles followed by translation to place the coordinate system’s origin along the insular axis.

**Results:** In a sample of 32 patients, the angle between the AC-PC line and the insular axis ranged from -17° to 17° in the axial plane (θ) and 31° to 69° in the sagittal plane (φ). In the resulting coordinate system, the insular axis defines z = 0 and the MCA turning point defines y = 0. We developed a custom, open-access MATLAB graphical interface that allows intuitive implementation of this system for both surgical planning and postoperative analysis; implanted electrodes, laser fiber position, and ablation geometry can each be localized within this common space. As a demonstration of its utility for pooling data across subjects, we applied the transformation to a previously acquired intracranial electrophysiology dataset and found that anatomically consistent, effector-specific motor representations emerged across 18 subjects once electrode positions were expressed in insular-specific coordinates.

**Conclusion:** As stereotactic surgery for insular targets becomes more common with expanding scientific inquiry, an insular-specific coordinate system may facilitate operative planning and functional mapping, and may help standardize outcome assessment across patients and institutions.

**SIGNIFICANCE STATEMENT:** The insular cortex represents an increasingly important surgical target for therapeutic interventions, yet substantial person-to-person anatomical variability hampers standardized targeting and outcome comparison. The insula is simultaneously the subject of expanding scientific inquiry — into interoception, pain, autonomic regulation, salience processing, and sensorimotor representation — much of it now pursued through intracranial recording and stimulation in humans, where cohorts are small, electrode sampling is idiosyncratic, and progress therefore depends on pooling data across patients in a frame that respects insular gyral architecture. We present “*Insulotaxy*,” a stereotactic coordinate system built from consistent, easily identifiable local anatomical landmarks that accounts for the insula’s unique rotational relationship to standard brain coordinates. An open-source MATLAB tool transforms imaging into insular-specific coordinates, facilitating surgical planning for ablation and electrode placement while enabling standardized outcome reporting across institutions. By providing locally anchored, anatomically aligned coordinates rather than relying on whole-brain registration, this framework addresses a practical gap in functional neurosurgery and lays a foundation for pooling clinical and electrophysiological data as insular interventions become more prevalent.

## INTRODUCTION

The insular cortex has emerged as a therapeutic target for drug-resistant epilepsy, chronic pain, and neuropsychiatric disorders, owing to its central role in interoceptive processing and its extensive connections with limbic, somatosensory, and autonomic networks.^1^ Recent advances in stereotactic technique - particularly laser interstitial thermal therapy (LITT)^2^ stereo- electroencephalography (SEEG)-guided radiofrequency thermocoagulation^3^ and responsive neurostimulation (RNS)^4^ - have made this previously inaccessible region approachable with markedly reduced morbidity compared with open resection **[Figure 1]**.^5,6^ However, accurate target localization across patients remains challenging because of the insula’s complex three-dimensional structure and its deep position within the Sylvian fissure. Standard volume-based registration algorithms often fail to adequately align insular landmarks; even the best-performing methods achieve only modest performance (Dice coefficients 0.40-0.74) for insular structures.^7^ Critically, these approaches do not account for the relative internal rotation of the insular gyri with respect to standard stereotactic spaces, introducing spatial errors that compromise cross-patient comparison of outcomes.

**Figure 1.**
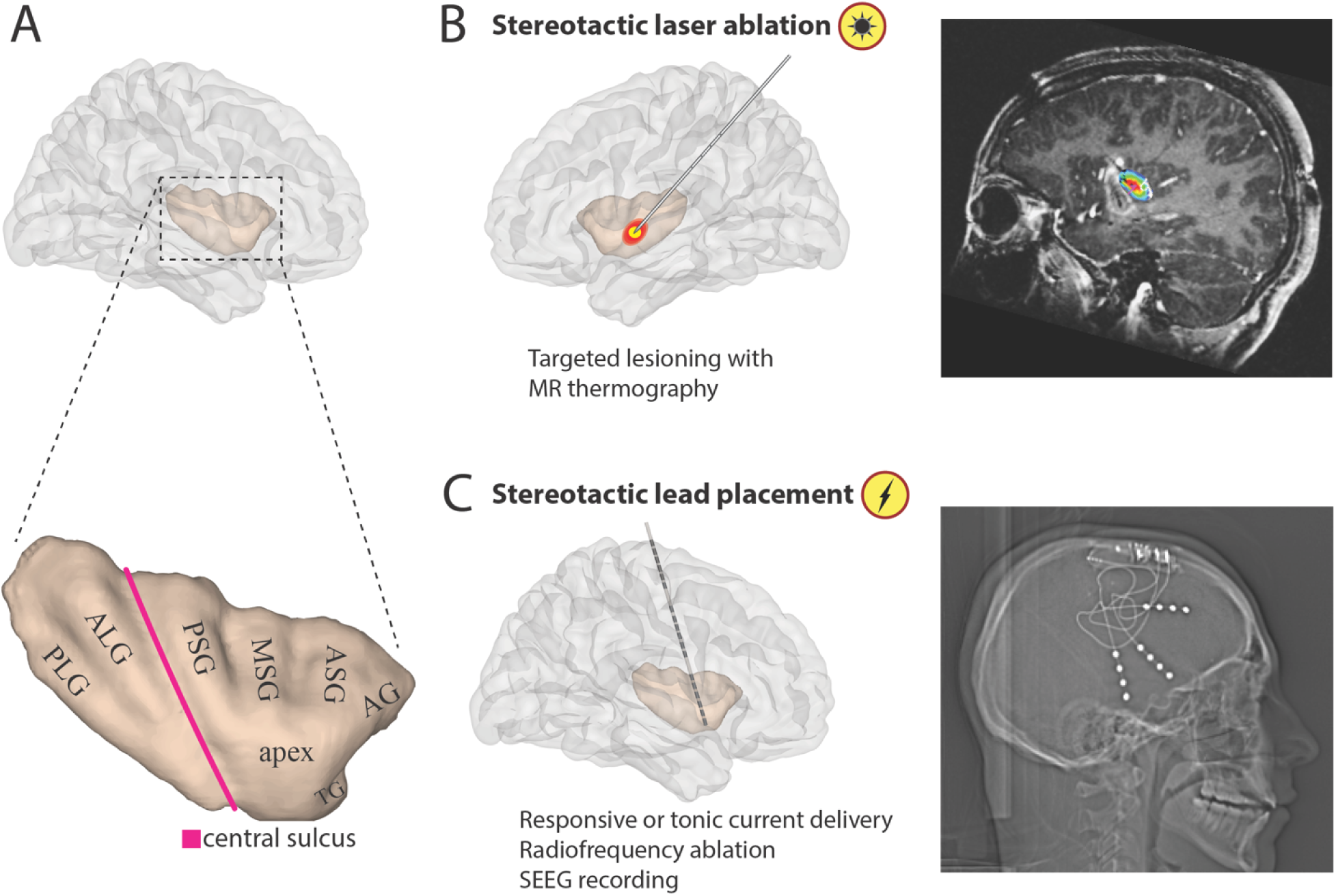
Insular macro-anatomy and stereotactic surgical approaches. Top panel: **(A)** Demonstration of insular macro-anatomy. The central sulcus of the insula (pink line) divides it into anterior and posterior regions. The anterior region consists of three short gyri (anterior/ASG, middle/MSG, posterior/PSG) that converge at the insular apex; the posterior region consists of two long gyri (anterior/ALG, posterior/PLG). An accessory gyrus (AG), anterior to the ASG, and a transverse gyrus (TG) on the basal surface continuous with the orbitofrontal cortex, are present in a subset of hemispheres. Bottom panel: Stereotactic approaches to the insular cortex have transformed the surgical treatment of epileptic and neuropsychiatric conditions, including **(B)** fiber-optic probes for laser ablation and **(C)** electrodes for recording of seizure activity, radiofrequency thermocoagulation, or tonic/responsive electrical stimulation.

Region-specific coordinate systems have proven valuable for other deep structures, with the AC-PC system providing standardized targeting for subcortical nuclei^8^ and specialized hippocampal coordinates enabling precise mesial temporal localization.^9^ These systems derive their power from anatomical landmarks specific to the region of interest rather than whole-brain transformations. No analogous coordinate system has yet been established for the insular cortex, and the consequences of this gap are well documented: Kang et al^7^ found that widely used nonlinear registration algorithms performed inconsistently when applied to the insula and its individual gyri, while Callaert et al^8^ demonstrated that the choice of segmentation and normalization pipeline most strongly affects morphometric outcomes in cortex adjacent to major sulci and fissures - a description that applies directly to the insula’s position deep within the Sylvian fissure.^10^

To address this gap, we present “*Insulotaxy*,” a stereotactic coordinate system designed specifically for the insular cortex and based on five reproducible anatomical landmarks: four points defining the central sulcus of the insula and one inferior point at the middle cerebral artery (MCA) bifurcation (the limen insulae). The system calculates rotational transformations in the axial and sagittal planes relative to standard AC-PC coordinates, accommodating the insula’s variable orientation while preserving the computational simplicity of a local, linear transformation. We have developed an open-source MATLAB graphical user interface that transforms pre- and postoperative imaging into insular-specific coordinates. This paper presents the mathematical framework underlying Insulotaxy, characterizes its application across a patient cohort, and demonstrates its practical value for surgical planning, group-level electrophysiological mapping, and standardized outcome assessment. By providing anatomically meaningful coordinates aligned with insular gyral architecture, Insulotaxy addresses a critical need for comparing insular interventions across patients and institutions.

## MATERIALS AND METHODS

### Overview of insular macro-anatomy

The human insula, situated within the Sylvian fissure, is pyramidal in shape once the overlying fronto-parietal-temporal opercula are reflected.^11^ It is clearly demarcated from the surrounding cortex by the anterior, superior, and inferior peri-insular sulci and is divided by the central insular sulcus into a larger anterior and a smaller posterior insula **[Figure 1]**. The central insular sulcus follows an oblique trajectory similar to that of the central (Rolandic) sulcus and represents the insula’s main and deepest sulcus.^12^ The anterior insula contains three principal short gyri (anterior, middle, and posterior), together with an accessory gyrus (present in 82% of hemispheres) and a transverse gyrus (present in 86%).^12^ The posterior insula comprises the anterior and posterior long gyri. The limen insulae forms a basolateral threshold connecting the insular cortex to the anterior perforated substance and corresponds anatomically to the site where the middle cerebral artery bifurcates or trifurcates into its major branches.

### Overview of the Insular Coordinate System

Our insular stereotactic coordinate system is implemented as a two-stage transformation: the brain volume is first aligned to standard anterior commissure-posterior commissure (AC-PC) space, and then subjected to a region-specific transformation into insular coordinates. This sequential approach preserves compatibility with existing neuroimaging workflows while providing the additional domain-specific relevant information required for insular targeting.

### Identification of Reference Landmarks

We first identify the landmarks required to transform the image into AC-PC space. The AC and PC are localized in standard fashion on the MRI, and three additional midline points are identified to define the midline plane; a rotation matrix is then calculated to align the structural MRI with the AC-PC coordinate system. We next identify the insular-specific landmarks. The central sulcus of the insula provides an intuitive axis that is identifiable across imaging modalities and is approximated by selecting multiple points along its inferior-to-superior extent. The limen insulae serves as the anterior landmark, marking the junction between the insula and the orbital surface of the frontal lobe.

### Plane of Symmetry and Axial Definition

To establish the coordinate system’s reference planes, we first define the plane of midline symmetry: the AC, PC, and additional midline landmarks are entered into a principal component analysis (PCA) to determine the normal vector to this plane. The insular central sulcus axis, defined by the landmark points above, forms the primary axis of the coordinate system, and the ventral-dorsal vector is defined as the cross product of the midline-plane normal vector and the insular axis vector **[Figure 2]**.

**Figure 2.**
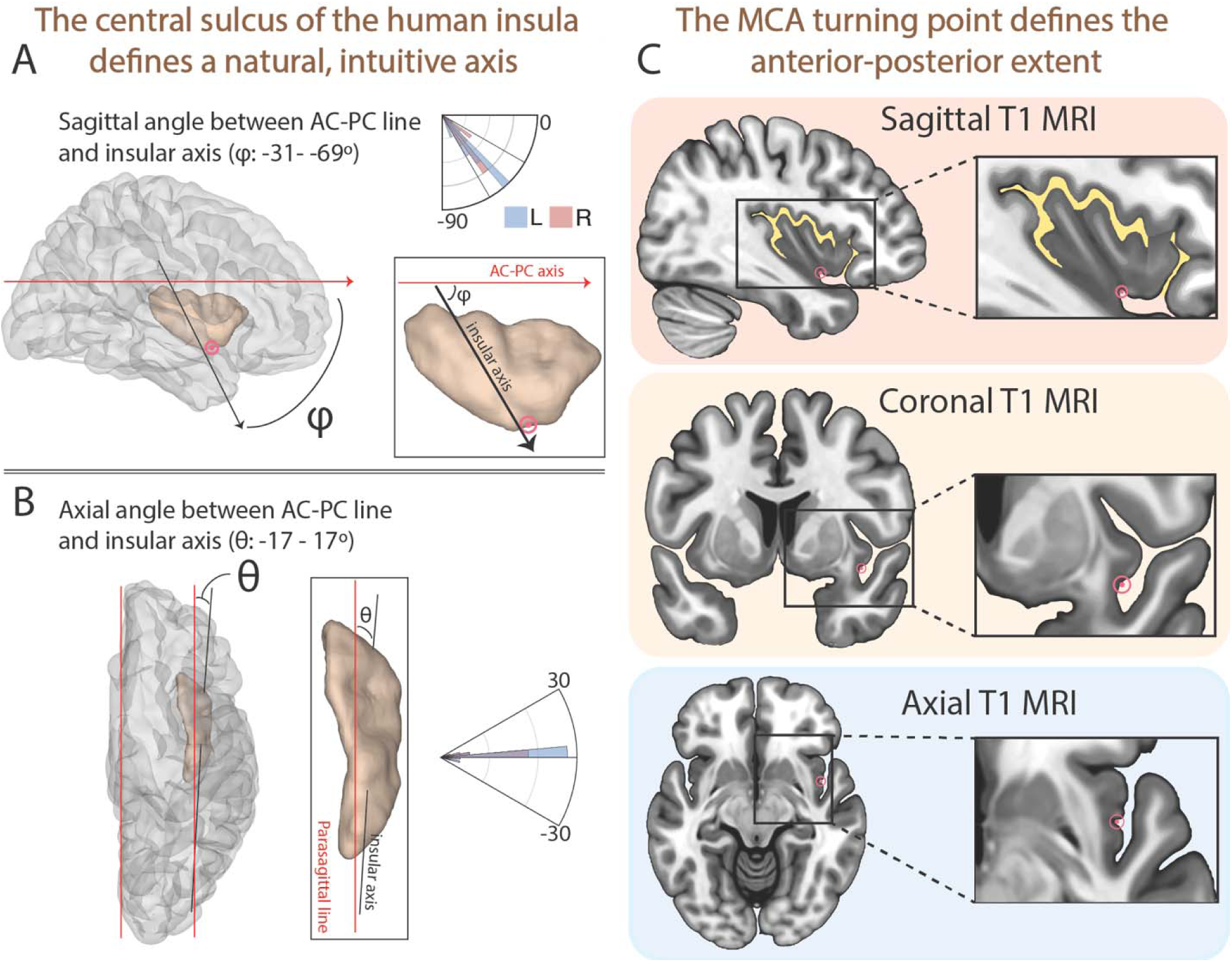
Angles defined by the insular axis. The insular axis, defined by the central insular sulcus, forms **(A)** a sagittal angle φ and **(B)** an axial angle θ with the AC-PC line; these angles are used to rotate each individual brain from AC-PC space into insular stereotactic space. **(C)** An indentation is typically observed at the inferior corner of the insula, where the middle cerebral artery (MCA) takes a steep turn before its bifurcation/trifurcation. This point was selected to define the anterior-posterior extent of the coordinate system (y = 0).

### Volumetric Rotation

Transformation into insular stereotactic space requires three sequential volumetric rotations: (1) the brain is rotated into alignment with standard AC-PC space; (2) the volume is rotated about the z-axis to align with the projection of the insular central sulcus onto the axial plane; and (3) the volume is rotated to align with the angle between the AC-PC line and the insular central sulcus axis in the sagittal plane. The complete transformation matrix is the multiplicative product of these individual rotation matrices, allowing the entire operation to be applied as a single affine transformation.

### Center of Coordinate System

Following rotation, the volume is translated so that the coordinate system’s origin lies at the intersection of the insular central sulcus and the limen insulae. This origin is anatomically consistent, readily identifiable across subjects, and relatively stable even in the presence of pathology or atrophy.

### Software Implementation

We developed a MATLAB-based software package, “*Insulotaxy*,” to implement this coordinate system through a graphical user interface for landmark selection, image registration, and coordinate transformation **[Figure 3]**. Imaging data are converted from DICOM to NIfTI format, after which the relevant landmarks are manually selected on preoperative T1-weighted MRI. The software then performs the affine transformation and reslices the volume into 1 mm^3^ insular stereotactic space. Additional imaging series are co-registered and resliced into this reference space using normalized mutual information, permitting multimodal integration of pre- and postprocedural images for precise localization of structures, trajectories, or regions of interest.

**Figure 3.**
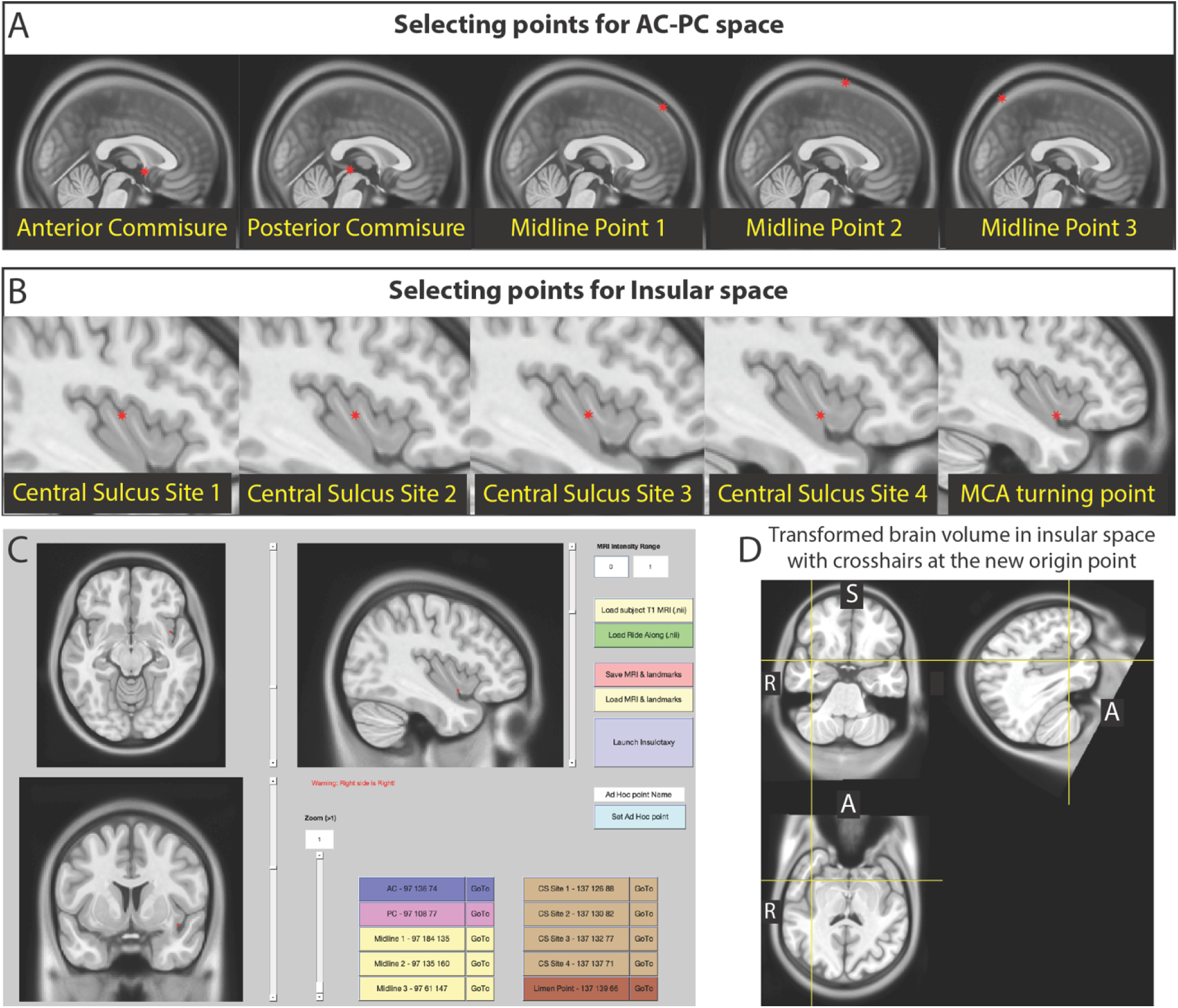
Graphical user interface for transformation into insular stereotactic space. **(A)** The user first identifies five points required to rotate the volume into AC-PC stereotactic space, **(B)** followed by five points required to rotate into insular space. **(C)** Example transformation of the right insula in the MNI-152 2009c nonlinear asymmetric, skull-stripped template brain. **(D)** Once the transformation is complete, the brain volume is rotated into the new coordinate space, which can then be used to visualize group-level data (see Figure 4).

### Practical Implementation for Surgical Planning

For surgical planning, the necessary transformation is calculated from the preoperative imaging, and the intended trajectory is defined within insular coordinates before being transformed back into native image space for surgical navigation. This preserves standardized targeting strategies while accommodating individual anatomical variation.

The same transformation can be applied after the procedure to map implanted electrodes, ablation zones, or other interventions into standardized insular space, enabling quantitative assessment of targeting accuracy and pooled analysis across patients.

### Application to Group-Level Electrophysiological Data

As a demonstration of the coordinate system’s utility for pooling data across subjects, we applied the *Insulotaxy* transformation to a previously acquired intracranial electrophysiology dataset from our group (Institutional IRB no. 15-006530).^13^ The cohort consisted of 18 subjects with drug-resistant epilepsy undergoing SEEG recording as part of their clinical workup. The postimplant CT scan was co-registered to the preimplant MRI using SPM12. Subjects performed a visually cued, self-paced hand, tongue, and foot motor execution task and the degree of neural activation was quantified using high-frequency broadband activity (65-115 Hz) They underwent preoperative 3T T1 MRI with and without intravenous gadolinium contrast using a standardized protocol on a Siemens MRI scanner. No patient in this cohort exhibited gross distortion of insular anatomy (e.g., lesion or encephalomalacia) that would confound assessment of the coordinate system. All experiments were performed in the epilepsy monitoring unit or the

pediatric intensive care unit. Individual electrode coordinates from all 18 subjects (196 bipolar channels) were transformed into insular stereotactic space using the methods described above and projected onto MNI-152 template brain for group-level visualization **[Figure 4]**. This analysis serves as an external, functional validation of the coordinate system’s capacity to preserve biologically meaningful spatial relationships when data are pooled across a large, anatomically heterogeneous cohort.

**Figure 4.**
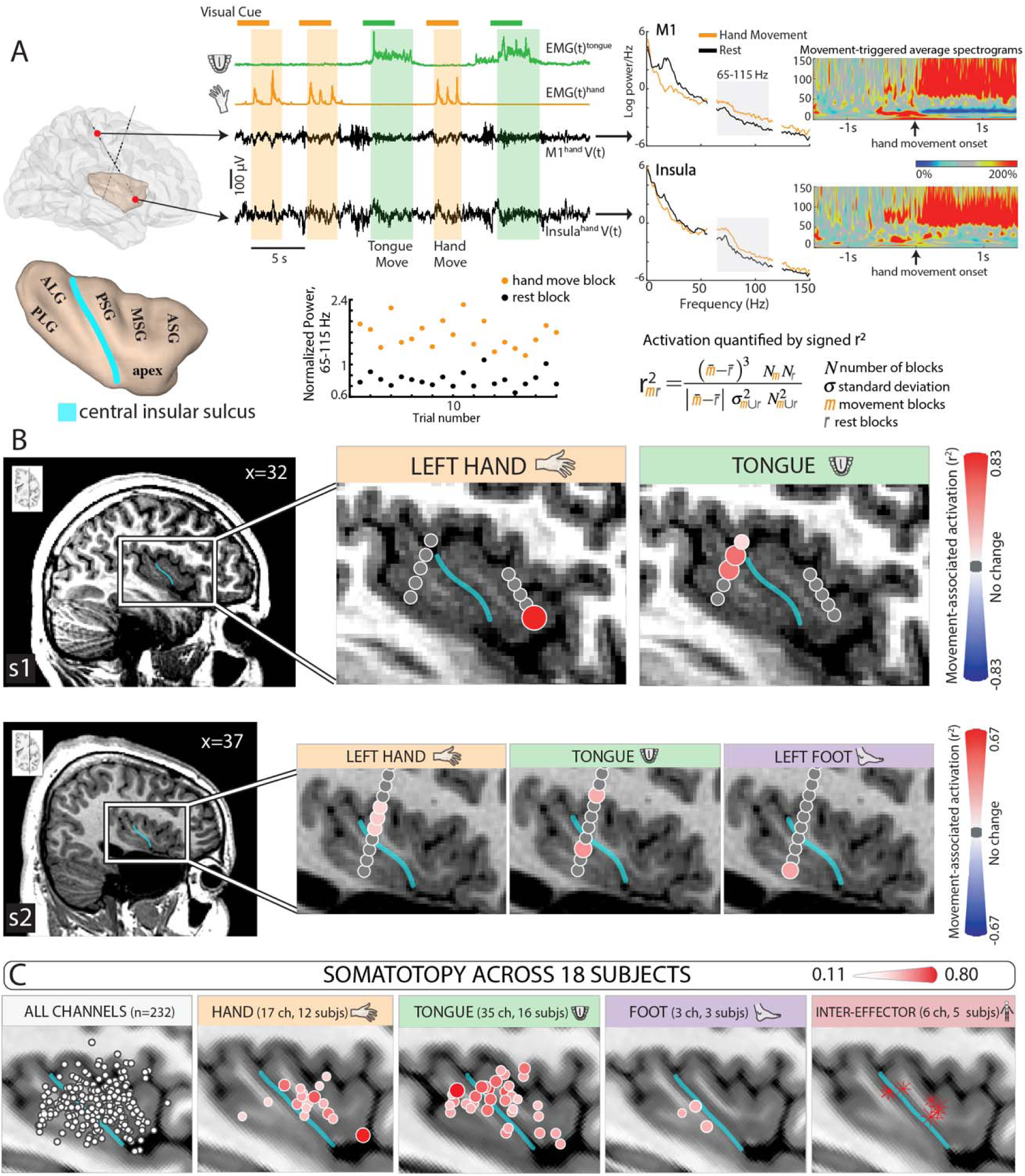
Application of the insular stereotactic coordinate system to pooled electrophysiological data. **(A)** Representative single-subject example of the movement-activation analysis used to generate the group map in (C): for each stereo-EEG channel, the bipolar-referenced broadband signal (65-115 Hz) is compared between movement and rest blocks, and the degree of activation is quantified with a signed-rank correlation coefficient (r2). The central insular sulcus (blue line) separates the anterior short gyri (anterior/ASG, middle/MSG, posterior/PSG), which converge at the insular apex, from the posterior long gyri (anterior/ALG, posterior/PLG). **(B)** Subject-level r2 activation maps for hand (orange), tongue (green), and foot (purple) movement in two representative subjects, displayed in the sagittal plane; each map is scaled to that subject’s maximum r2 value. **(C)** After transforming each subject’s electrode coordinates into insular stereotactic space using the method described here, all intrainsular contacts (232 bipolar channels; 18 subjects) are displayed in a common reference frame on the MNI-152 template. Hand-tuned sites (orange) cluster along the ventral middle and posterior short gyri bilaterally, and tongue-tuned sites (green) cluster along the dorsal posterior short gyrus and anterior long gyrus - an anatomically consistent, effector- specific grouping that emerges only once electrode positions are expressed in insular-specific, rather than whole- brain, coordinates. Adapted from Kerezoudis et al (license to reuse under under Creative Commons Attribution- NonCommercial-NoDerivatives License 4.0 (CC BY-NC-ND).^13^

### Data Availability

All anonymized data recorded necessary to interpret, verify and extend the research is publicly available at Open Science Framework (at https://osf.io/bmdy6/overview).

### Code Availability

All code necessary to reproduce these findings will be made publicly available on GitHub upon publication of the manuscript (at https://github.com/Kerezoudis). A user manual is also available there. All analyses were conducted in MATLAB (R2024a, MathWorks, Natick, MA, USA). Figures were produced in MATLAB and edited in Adobe Illustrator 2024 (Adobe, San Jose, CA, USA).

## RESULTS

Using our open-access, custom MATLAB software, we identified the five anatomical landmarks required for registration and calculated the rotational angles between the AC-PC line and the insular axis, as defined by the central insular sulcus **[Figure 2]**. The sagittal angle (φ) between the AC-PC line and the insular central sulcus axis measured -47.7° ± 7.9° (range: -57.2° to -31.4°) in the right hemisphere (n = 17) and -51.4° ± 7.8° (range: -69° to -37.2°) in the left hemisphere (n = 22), with no significant difference between hemispheres (p = 0.15). The axial angle (θ) between a parasagittal line (parallel to the AC-PC line) and the insular central sulcus axis measured 3.5° ± 6.2° (range: -10.1° to 16.7°) in the right hemisphere and -0.8° ± 6.4° (range: -17.3° to 9.7°) in the left hemisphere, a difference that reached statistical significance (p = 0.04). **Figure 2** further illustrates the limen insulae and its consistent identifiability as the anterior/inferior landmark of the coordinate system.

**Figure 3** demonstrates the Insulotaxy graphical user interface, illustrating how users select the requisite landmarks, generate the affine transformation, and visualize the resulting insular-space volume. This interface allows rapid, intuitive implementation of the coordinate system and facilitates both prospective surgical planning and retrospective postoperative analysis.

As proof-of-concept for the coordinate system’s capacity to support cross-subject data pooling, we applied the Insulotaxy transformation to a previously acquired intracranial electrophysiology dataset examining movement-related high-frequency broadband activity (65- 115 Hz) in the insula during a visually cued hand, tongue, and foot motor task.^13,14^ Individual insular electrode coordinates from 18 patients (196 bipolar channels) were each transformed into insular stereotactic space and projected onto a common brain surface for group-level visualization (**Figure 4**). Despite substantial inter-individual variation in insular gyral anatomy and heterogeneous electrode sampling across patients, this approach revealed anatomically consistent, effector-specific clustering across the cohort: hand-tuned sites concentrated along the ventral middle and posterior short gyri, while tongue-tuned sites clustered along the dorsal posterior short gyrus and long gyri. This finding demonstrates the practical utility of the coordinate system for aggregating electrophysiological - and, by extension, other multimodal - data across a large and anatomically heterogeneous patient cohort, an analysis that would otherwise be confounded by the misalignment inherent to whole-brain registration methods.

## DISCUSSION

### Research applications

For research applications, Insulotaxy addresses a specific methodological gap in functional neuroimaging of the insula. Standard nonlinear volumetric approaches, such as SPM12’s unified segmentation, do not take into account the intrinsic geometry of insular gyri, whereas surface-based methods, such as Freesurfer, are labor-intensive and can take up to 24 hours to complete.^15,16^ Surface methods also inflate the cortical ribbon and morph it onto a sphere, a mapping that minimizes but cannot eliminate metric distortion, so distances and angles are least faithfully preserved in deeply folded, anatomically variable cortex.^17^ They are further vulnerable to failure when surface artifact corrupts the pial or white matter boundary, as with a ventriculoperitoneal shunt or prior craniotomy, whereas a landmark-based rigid transformation is robust to such artifact. The *Insulotaxy* framework therefore balances anatomical precision and computational simplicity for insular localization. Constraining the transformation to five reproducible landmarks achieves alignment of functionally relevant insular structures while preserving local geometric integrity. This allows to easily capture fundamental gradients in insular functional organization, which includes anterior-to-posterior transitions from socio- emotional to sensorimotor processing and dorsal-to-ventral gradients in interoceptive representation.^18,19^ We demonstrate this capacity directly in the present study: applying Insulotaxy to a previously acquired electrophysiological dataset from 18 subjects reveals consistent, effector-specific somatotopic clustering that would otherwise be obscured by the anatomical misalignment inherent to whole-brain registration. The same framework could standardize how white matter pathways are referenced in insular connectivity atlases and support meta-analyses of diffusion tractography studies. As understanding of insular function expands across cognitive, affective, and interoceptive domains, *Insulotaxy* offers a standardized framework for integrating findings across modalities and translating them into clinical application.

### Clinical applications

Clinically, Insulotaxy addresses challenges intrinsic to surgical approaches to the insula, which are complicated by its deep location, rich vascularization, and proximity to critical white matter tracts. The system provides a standardized framework for preoperative planning that respects the insula’s natural anatomy, enables precise documentation of SEEG contacts relative to insular gyral anatomy, and allows LITT ablation volumes and burn geometry to be recorded in standardized coordinates for comparison with seizure outcome across patients. Mapping prior interventional outcomes into a common reference space also facilitates evidence-based targeting and probabilistic mapping of therapeutic efficacy and complication risk — an increasingly relevant capability as RNS and other minimally invasive approaches expand the range of insular targets.

### Limitations

Several limitations merit consideration. The system requires manual identification of five anatomical landmarks, introducing rater-dependent variability; automated or semi-automated landmark detection trained on large annotated datasets is a natural next step. A linear transformation may also not fully capture anatomical variability in patients with atypical insular morphology from developmental variation, mass effect, or prior surgery, and prospective studies are needed to demonstrate improved targeting accuracy and clinical outcomes relative to conventional approaches.

*Insulotaxy* is also not intended as a universal replacement for whole-brain registration, and **Table 1** summarizes where each approach is best suited. Because the transformation is local to the insula, it is a poor fit for questions that require whole-brain correspondence: population-level voxel-based morphometry, integration with probabilistic cytoarchitectonic atlases, and surface-based analyses spanning the full cortical mantle remain better served by volumetric or surface registration, which also offer fully automated, higher-throughput reproducibility across large cohorts than a landmark-based method. Insulotaxy is best reserved for questions centered on the insula itself — stereotactic surgical planning, SEEG/LITT trajectory and electrode documentation, and pooling of insular electrophysiological or imaging data across patients — where preserved local geometry and intraoperative speed outweigh the value of whole-brain correspondence.

**Table 1.** Comparison of Registration Approaches for Insular Cortex Analysis & Targeting.

| Characteristic | Insulotaxy | Volumetric Registration (SPM12/ANTs) | Surface Registration (FreeSurfer) |
| --- | --- | --- | --- |
| **Technical Approach** | Affine transformation using 5 insular landmarks | Nonlinear voxel-wise warping to MNI template | Surface-based spherical registration |
| **Processing Time** | <5 minutes | 10-20 minutes | 6-24 hours |
| **Manual Input** | 5 landmarks | Minimal (automated) | None (automated) |
| **Insular Alignment** | Excellent (designed for insula) | Modest (Dice: 0.40-0.74) | Variable (struggles with buried cortex) |
| **Local Geometry Preservation** | Excellent | Good (nonlinear warping) | Very good (cortical surface only) |
| **Surgical Planning** | Excellent | Limited | Poor |
| **Intraoperative Feasibility** | Yes | No | No |
| **Coordinate Interpretation** | Intuitive (axis alignment) | Abstract (mm from AC) | Abstract (vertex-based) |
| **Reproducibility** | Very good (rater-dependent) | Excellent (automated) | Excellent (automated) |
| **Best Applications** | Stereotactic surgery, SEEG/LITT planning, clinical trials | Whole-brain analysis, MNI integration | Cortical morphometry, surface fMRI |
| **Primary Limitation** | Requires manual landmarks | Suboptimal insular specificity | Not suitable for stereotaxy |

**Abbreviations:** LITT, laser interstitial thermal therapy; SEEG, stereoelectroencephalography; fMRI, functional magnetic resonance imaging; MNI, Montreal Neurological Institute; AC, anterior commissure

## CONCLUSION

As stereotactic and ablative approaches to the insula become more common, *Insulotaxy* offers an anatomically grounded alternative to whole-brain registration for surgical planning, electrode and lesion localization, and the pooling of clinical and electrophysiological data across patients. It could help standardize how insular interventions are planned, reported, and compared across institutions — supporting evidence-based refinement of surgical technique for patients with drug-resistant epilepsy, chronic pain, and other conditions amenable to insular intervention.

## Disclosures

We are grateful to the patients who volunteered their time to participate in this research, to B. Wessel, C. Nelson and the staff at St. Mary’s Hospital. This work was supported by the NIH-NCATS CTSA KL2 TR002379 (KJM), the Foundation for OCD Research (KJM), and by the Brain Research Foundation with a Fay/Frank Seed Grant (KJM), NIH U01-NS128612 (KJM, GAW), and NIH R01-MH122258 (DH). The contents of this manuscript are solely the responsibility of the authors and do not necessarily represent the official views of the NIH.

GAW reports licensed intellectual property related to neurotechnologies with Cadence Neuroscience and NeuroOne Medical Technologies. He serves on Scientific Advisory Boards for NeuroPace, LivaNova, NeuroOne Medical Technologies, Cadence Neuroscience, and UNEEG Medical. He has received research funding from Medtronic, UNEEG Medical, Cadence Neuroscience, and Modulight Inc. These relationships are outside the submitted work unless otherwise noted. Our funders did not play a role in study design, data collection and analysis, decision to publish, or preparation of the manuscript.

## Author contributions

**P.K.** - Conceptualization and design, data collection, analysis and drafting of manuscript

**M.A.J.** - Conceptualization and design, data collection, drafting of manuscript

**B.T.K.** - Study supervision, reviewing and revising original draft

**G.A.W.** - Study supervision, reviewing and revising original draft

**N.F.I.** - Study supervision, reviewing and revising original draft

**J.V.G.** - Study supervision, reviewing and revising original draft

**K.J.M.** - Conceptualization and design, study supervision, data collection, reviewing and revising original draft

## Notes

https://osf.io/bmdy6/overview

### Competing Interest Statement

The authors have declared no competing interest.

